# Enteric pharmacokinetics of monomeric and multimeric camelid nanobody single-domain antibodies

**DOI:** 10.1101/2023.05.15.540785

**Authors:** Michelle Debatis, Hillary Danz, Jacqueline M. Tremblay, Kimberly Gaspie, Raymond K. Kudej, Vladimir Vigdorovich, Noah Sather, Justyna J. Jaskiewicz, Saul Tzipori, Charles B. Shoemaker

## Abstract

Single-domain antibodies (sdAbs) derived from *Camelidae* heavy-chain-only antibodies (also called nanobodies or VHHs) have advantages over conventional antibodies in terms of their small size and stability to pH and temperature extremes, their ability to express well in microbial hosts, and to be functionally multimerized for enhanced properties. For these reasons, VHHs are showing promise as enteric disease therapeutics, yet little is known as to their pharmacokinetics (PK) within the digestive tract. To improve understanding of enteric VHH PK, we investigated the functional and structural stability of monomeric and multimeric camelid VHH-agents following *in vitro* incubation with intestinal extracts (chyme) from rabbits and pigs or fecal extracts from human sources, and *in vivo* in rabbits. The results showed that unstructured domains such as epitopic tags and flexible spacers composed of different amino acid sequences were rapidly degraded by enteric proteases while the functional core VHHs were much more stable to these treatments. Individual VHHs were widely variable in their functional stability to GI tract proteases. Some VHH-based agents which neutralize enteric Shiga toxin Stx2 displayed a functional stability to chyme incubations comparable to that of Stx2-neutralizing IgG and IgA mAbs, thus indicating that selected nanobodies can approach the functional stability of conventional immunoglobulins. Enteric PK data obtained from *in vitro* incubation studies were consistent with similar incubations performed *in vivo* in rabbit surgical gut loops. These findings have broad implications for enteric use of VHH-based agents, particularly VHH fusion proteins.

## Introduction

Enteric diseases substantially impact health in both human and veterinary medicine. While modern sanitation methods largely control enteric disease in the developed world, food-borne pathogens remain a major cause of disease and death in the developing world, particularly among children. Vaccines and drugs such as antibiotics and anthelmintics are available to prevent and/or treat many enteric diseases but are underutilized in the developing world due to factors such as cost, appearance of drug-resistant pathogens and inadequate medical infrastructure. Antibody therapeutics could also provide enteric disease benefits but are currently costlier and more challenging to distribute and administer than vaccines and small molecule drugs.

Single-domain antibodies (sdAbs) such as Camelid heavy-chain-only V_H_ (VHH, nanobody) domains offer the potential to overcome some of the obstacles to enteric immunotherapies. VHHs can be identified that bind to pathogen targets with similar affinities to conventional antibodies, yet possess advantages such as reduced production costs, improved stability to temperature and pH extremes, expression as multimers with enhanced properties, and amenability to gene therapies (recently reviewed by (1)). A number of reports have emerged in which VHH agents have been successfully employed as oral prophylactic agents to prevent rotavirus infections (2–4), ETEC (5), inflammatory bowel disease (6, 7) and pathology related to *Clostridium difficile* infection (8). To bypass potential damage to VHHs within the stomach, in some cases oral VHH delivery employed encapsulation in nanoparticles (7), delivery as fusions to IgA Fc domains within plant seeds (9) or expression by genetically modified blue-green algae (spirulina) (10) and probiotic bacteria such as *Lactobacillus* and *Lactococcus* (4, 6, 8, 11).

Despite the successes reported using oral therapeutic VHHs to treat enteric pathology, studies to evaluate the PK of VHHs within the GI tract are very limited. A major limitation to performing these studies is the inability to reliably detect functional VHHs that survive exposures to enteric proteases using routine immunological methods. This is because antibodies may falsely detect epitopes on partially degraded, non-functional VHHs or fail to detect surviving functional VHHs due to selective degradation of the epitopic tag used by the assay. For this reason, enteric studies on VHH PK are best performed by monitoring the survival of a specific VHH function in place of immunological methods. For example, Maffey et al (2) monitored rotavirus neutralization of VHHs following treatments with simulated stomach or intestinal fluids and showed significant degradation by pepsin but little degradation with trypsin. It is not clear though how accurately the simulated fluids represent the actual GI tract environment. In another study, VHH enteric PK was evaluated in monkeys given encapsulated VHHs (12) by assessing the ability of a protease-exposed VHH to compete with a mAb for binding its common epitope on human TNFα by ELISA. By this method, significant amounts of functional anti-human TNFα VHH were detected in the GI tract and feces of the monkeys, and later in human patient feces (7). Because it is not known at which times the VHHs were released by the protective nanoparticles, it is difficult to estimate the actual functional half-life of the released VHHs in the enteric proteolytic environment. Other studies have clearly demonstrated the survival of functional VHHs in the GI tract through demonstrations of protection from the enteric pathogen targeted by the VHHs following various oral treatment modalities (3–6, 8), but these studies were not accompanied by PK analyses.

In this manuscript, we employ extracts prepared from the upper intestine of rabbits and piglets (chyme), and human fecal extracts, to perform *in vitro* studies that characterize the stability of VHH-based agents to enteric proteases. The studies were done primarily employing functional assays to assess VHH monomer and heterodimer survival to enteric protease exposure. We developed and characterized a rabbit anti-VHH serum that we demonstrate to be capable of improved core VHH recognition in ELISA and western blot assays, thus permitting the acquisition of immunological data in support of the functional data. Finally, we performed rabbit enteric PK studies in live animals to seek *in vivo* validation of the *in vitro* approaches. The results clearly show that the enteric proteases rapidly degrade unstructured regions on the VHH-based agents, such as epitopic tags and different spacers sequences, while the core VHH function is far more resistant to the proteases. Different VHHs appear to have highly variable susceptibility to enteric protease inactivation. The results should provide guidance to researchers seeking to develop effective VHH-based therapeutics for treatments of enteric diseases.

## Results

### Human fecal and porcine intestinal extracts appear to preferentially cleave the spacers separating VHH multimers

Early western blot studies incubating intestinal and fecal extracts with VHH multimers had suggested that the proteins were preferentially cleaved within the spacer regions separating the VHH monomer components. A particularly clear example occurred when we incubated a heterohexameric agent consisting of VHHs linked by conventional (GGGGS)_N_ spacers (13) with dilute extracts (final 1:50 dilution) obtained from human feces or porcine intestine. As shown in **Fig 1**, western blots that recognize the epitopic E-tags which flank the six linked VHHs produced a similar partial digestion pattern when incubated with either human or porcine enteric proteases. The size of the products, multiples of ∼15 kDa, suggested that both extracts cleaved the heteromultimer within the spacers separating the 15 kDa VHH components.

**Fig 1.**
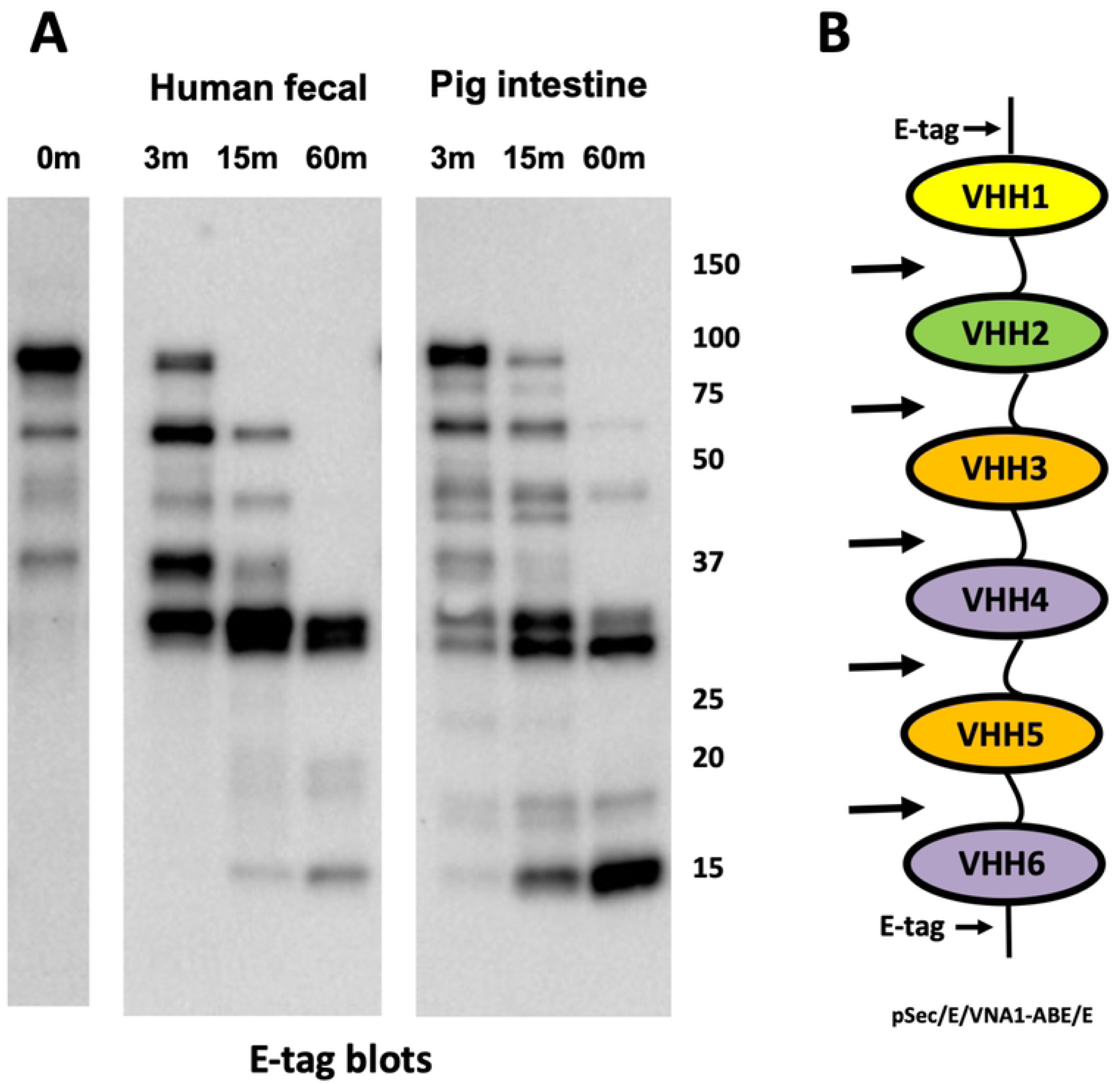
VHH heterohexamer degradation by intestinal proteases. (**A**) Western blot of a VHH heterohexameric protein (50 µg/ml) following incubation with extracts of human feces or porcine intestine. Intestinal extracts were added at a 1:50 dilution to serum-free conditioned medium of CHO cells transfected with the previously described VHH heterohexamer VNA1-ABE (13). Samples were removed following the indicated times of incubation (37°C) and quenched, and a western blot was performed detecting E-tag. (**B**) Diagram of VNA1-ABE indicating the positions of the E-tags which flank the VHH heterohexamer. The arrows indicate the approximate positions of the dominant intestinal protease cleavage sites within the protein.

### Core VHHs resist enteric protease degradation longer than peptide tags and spacers

To further characterize the degradation of VHH heteromultimers by enteric proteases, two different purified VHH heterodimers, each flanked by E-tag peptides and fused to thioredoxin (Trx) as shown in **Fig 2A**, were incubated at 37°C for 10 or 60 minutes with PBS or with extracts of human feces or porcine intestinal chyme (1:10 each). One of the heterodimers (Trx/E/ALcH7/JPUA5/E) consisted of two VHHs that potently inhibit the activity of Botulinum neurotoxin A (BoNT/A) light chain protease (LC/A) (14). The second heterodimer (Trx/E/JLEG10/JLEE9/E) consisted of two negative control VHHs binding a different protease (15). Aliquots of the incubation products were subjected to SDS-PAGE and blotted for products containing an E-tag. As shown in **Fig 2B**, the E-tags were rapidly lost after 10 minutes of incubation and became undetectable after 60’ incubation. Chyme proteases were inhibited with α1-antitrypsin (α1AT), and then the incubation products were assayed for LC/A inhibition (**Figs 2C and S1**). As a zinc protease, LC/A is not inhibited by α1AT. Surprisingly, all samples of the LC/A inhibitory VHH heterodimer retained substantial potency (i.e., preventing SNAP25 cleavage by LC/A) even after 60’ incubation with human fecal or pig intestinal extracts, and despite being virtually undetectable with anti-E-tag reagent on the western blot. Clearly, the flanking E-tag epitopes were becoming degraded or excised from the VHH heterodimers during enteric extract incubation long before the VHH agents lost their binding and inhibition activity. In other studies, we employed VHH agents fused to other epitope tags such as myc-tag, S-tag and hexahistidine tags, however all were rapidly degraded following similar incubations with GI extracts and thus poorly useful for PK studies (not shown). Our results strongly suggested that the highly-structured functional core VHHs are much more resistant to enteric proteases than the flanking epitopic tags. The poor correlation between VHH immunological detection and VHH function when monitoring enteric VHHs with conventional anti-tag antibody reagents highlights the substantial challenge of performing simple and quantitative enteric pharmacokinetics (PK) studies on therapeutic proteins.

**Fig 2.**
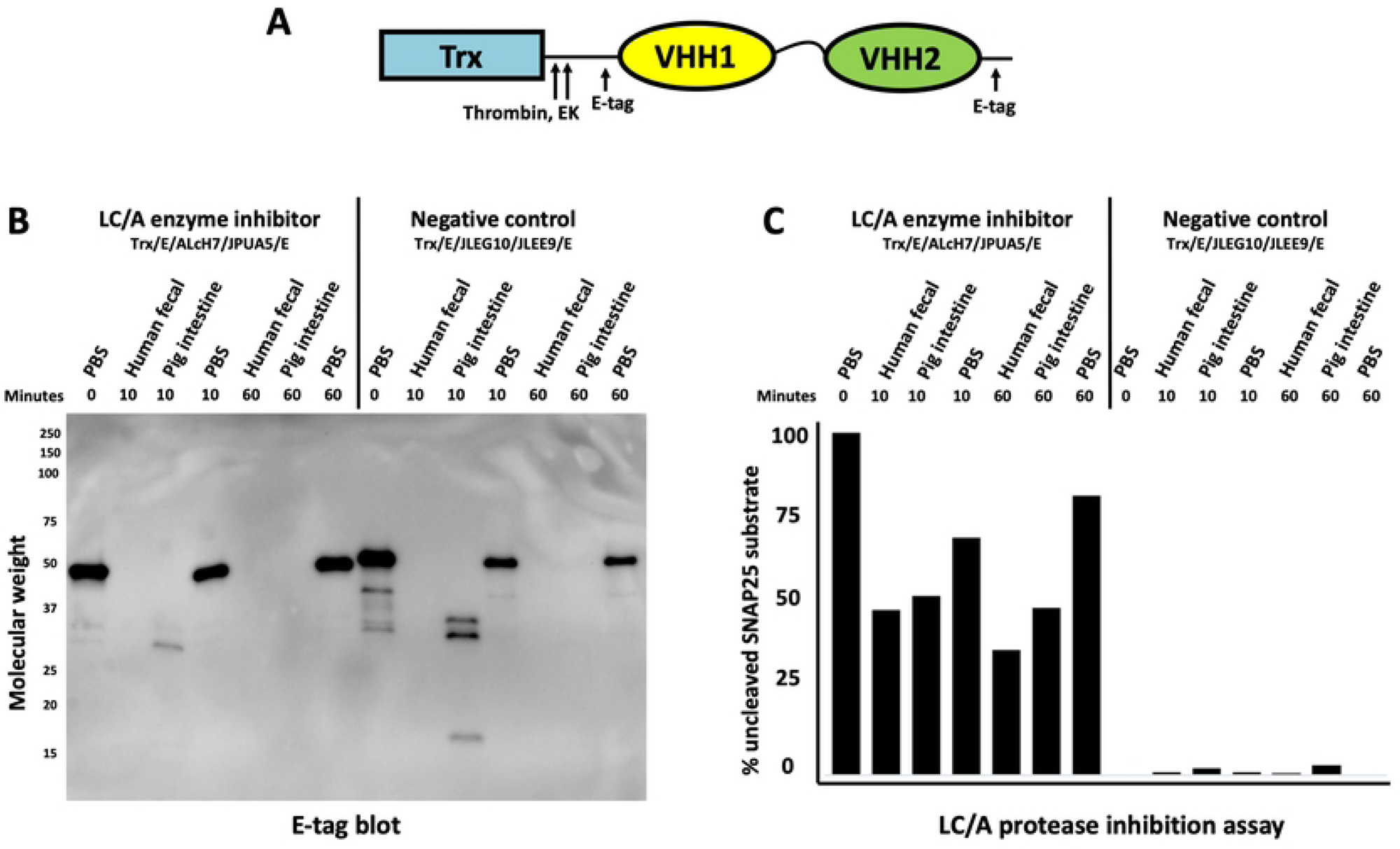
Intestinal proteases preferentially degrade epitope tags and VHH spacers before inactivating VHH function. (**A**) Diagram of a standard VHH heterodimer employed in these studies. The VHH heterodimers consisted of two different VHH components (VHH1, VHH2) separated by a spacer ((GGGGS)3), flanked by two E-tags and with an amino terminal *E. coli* thioredoxin (Trx) fusion partner for improved expression. These recombinant proteins derive from a modified pET32 expression vector and thus also contain both protease sensitive thrombin and enterokinase (EK) cleavage sites between the Trx and the VHH heterodimer as shown. Extracts were added at a 1:10 chyme dilution to two VHH heterodimer proteins (20 µg/ml). Samples were removed at indicated times of incubation (37°C) and quenched for later analysis. (**B**) Aliquot VHH heterodimer samples (100 µg/ml) incubated with chyme (1:10) for the indicated times were then added to SDS sample buffer and boiled. 10 µl aliquots were resolved by SDS-PAGE and a western blot was performed detecting E-tag. (**C**) Chyme exposed samples were adjusted to 100 µg/ml α1-antitrypsin (α1AT) to inhibit intestinal protease degradation and rapidly frozen. Later, BoNT/A protease (LC/A) was added to the thawed samples and assessed for their LC/A inhibitory activity. Raw data used for this assay is shown in **Fig S1**.

### Sensitive enteric protease cleavage sites within VHH heterodimers

We next sought to identify some specific sites in VHH agents which are preferentially cleaved by human and porcine enteric proteases. A series of studies were performed employing different VHH heterodimers. The purified VHH heterodimers were digested for various times with human fecal or porcine intestinal extracts to identify time points at which distinct digestion products became apparent. Sufficient protein was employed to permit identification of the amino terminal sequences of digestion products by Edman degradation methods. A summary of four cleavage sites identified after digestion is shown in **Fig 3** and derive from data shown in **Figs S2-S7**. All cleavage sites were found to occur very close to the amino terminus of one or both core VHHs. No clear sequence specificity was apparent, though all cleavage products contained a small amino acid (glycine or alanine) at the amino end. The results suggested that preferred intestinal protease cleavage sites are located in the likely unstructured spacer regions that immediately precede the component core VHHs. This specificity is suggestive of cleavage by a gastrointestinal serine protease such as neutrophil elastase (16). Preferential cleavage at these unstructured sequences is consistent with the findings shown above in which substantial VHH function remains intact well after the epitopic tags have been removed by intestinal protease action.

**Fig 3.**
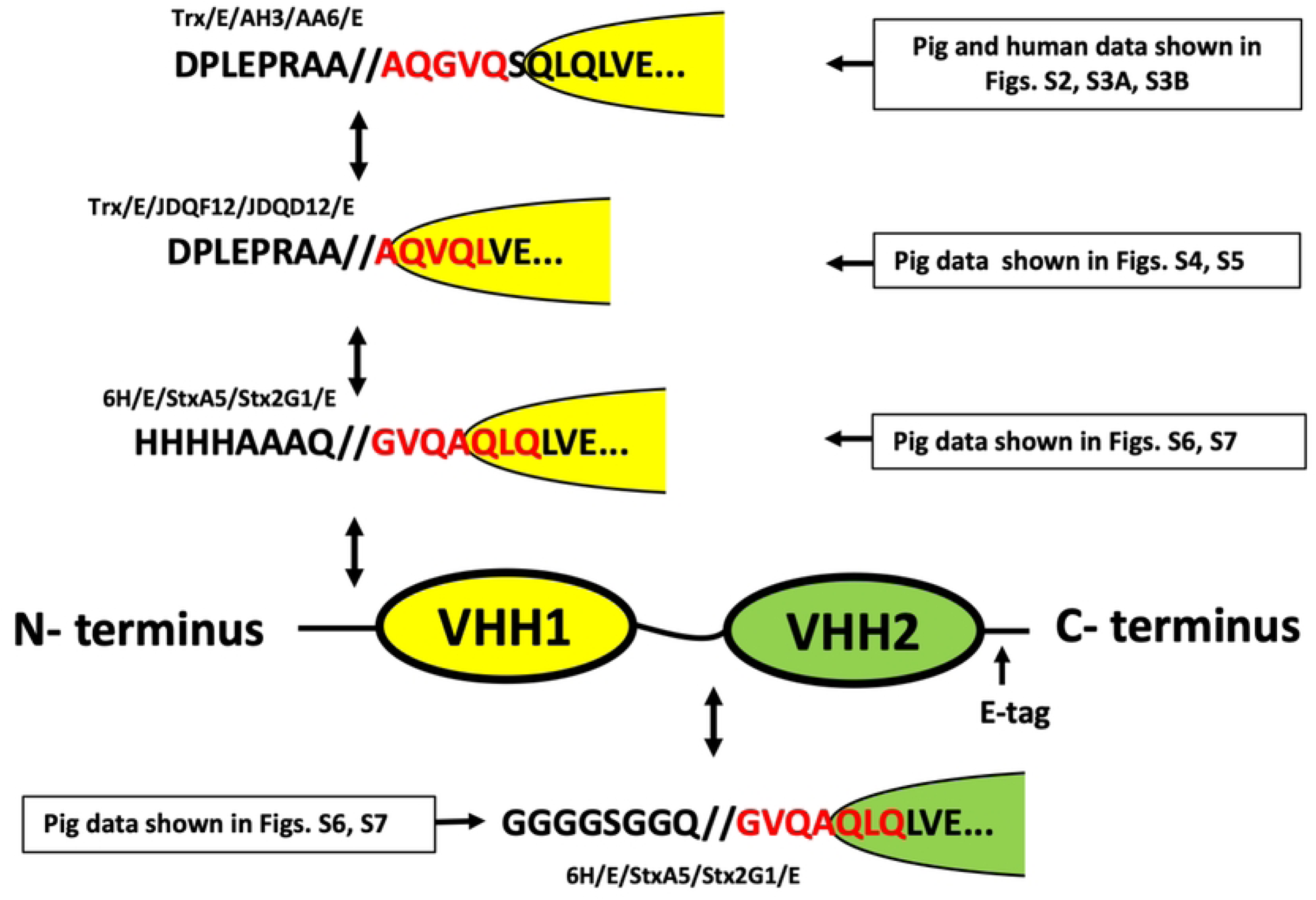
Preferred intestinal protease cleavage sites identified in multimeric VHH spacer and flanking regions. Shown are the amino acid sequences surrounding four intestinal protease cleavage sites identified by Edman sequencing (depicted in red) of purified intestinal protease degradation products. Arrows indicate the positions within a VHH heterodimer diagram at which the intestinal protease cleavage sites are found to have occurred. Also indicated are the supplemental figures (**Fig S2-S7**) that show the raw data from the Edman degradation sequencing employed for each of the cleavage site identifications.

### Development and testing of anti-VHH serum for enteric PK studies

After determining that immunologic detection of VHHs using conventional anti-epitopic tag reagents was unreliable following VHH exposures to intestinal proteases (**Fig 2**), we sought to identify an immunological reagent capable of detecting the functional VHH cores to facilitate more robust enteric PK studies. We initially immunized rabbits with a mixture of 13 different native VHH proteins in a non-denaturing aqueous alum adjuvant. Each of the VHHs in the pool also contained ∼50 pET32 vector-encoded, amino-terminal amino acids consisting of several purification epitopic tags and a short C-terminal myc tag (Table S1). Due to early unsatisfactory results (e.g., shown below) with the resulting immune sera (called ‘rabbit anti-native VHH sera’), we immunized a second pair of rabbits employing the same VHHs as before, but 50% of the VHH pool was removed, partially denatured (reduction and boiling) and recombined. The partially denatured pool was then formulated in an oil-based Freund’s adjuvant for all immunizations. The resulting immune sera was called ‘rabbit anti-denatured VHH sera’.

The two different rabbit anti-VHH sera preparations were tested for recognition of an Stx2-neutralizing VHH heterodimer (**Fig 4A**) sampled after various times of incubation (30m, 3h, o/n) with porcine intestinal chyme (1:20). In one analysis, samples were resolved by SDS-PAGE and detected by western blotting. As shown in **Fig 4B**, sera derived from rabbits immunized with anti-denatured VHHs contained antibodies that were capable of recognizing distinct peptides that preferentially survived long term intestinal protease incubations, such as those observed by Coomassie staining in **Figs S2, S4 and S6**. The smallest band recognized was ∼14 kDa, the predicted size of core VHHs. In contrast, rabbit anti-native VHH sera poorly recognized the VHH heterodimer or the VHH monomers after only 30 minutes exposure to intestinal extract, thus displaying similar sensitivity to intestinal proteases as previously found with anti-E-tag reagent (Fig. 2). This result suggests that the rabbit anti-native VHH sera almost exclusively recognizes the short unstructured amino and carboxy terminal peptides, such as the epitopic tags, common to all of the recombinant VHHs used for rabbit immunization.

**Fig 4.**
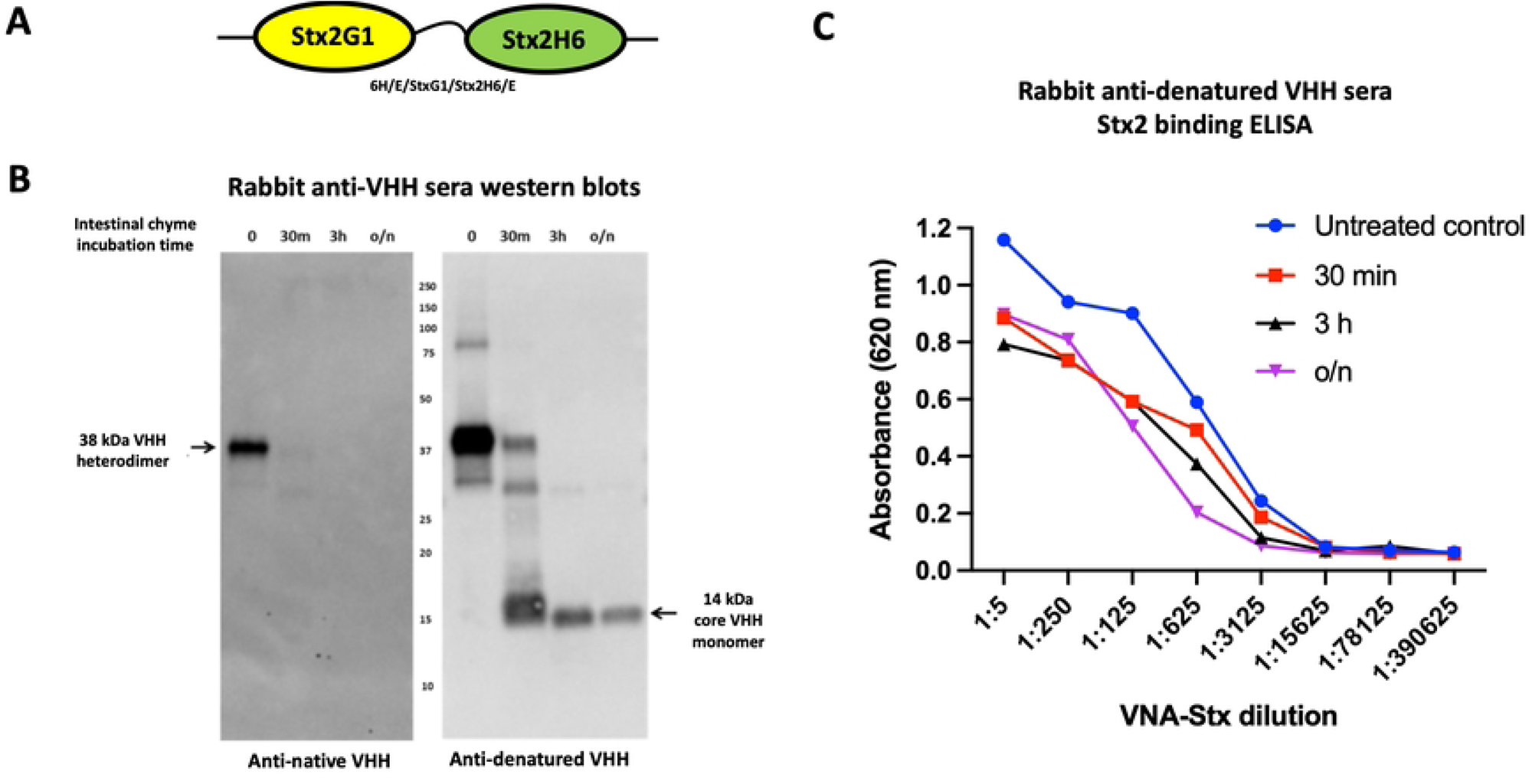
Characterization of sera from rabbits immunized with a VHH pool for their recognition of ‘core’ VHHs. (**A**) Diagram of a VHH heterodimer (6H/E/StxG1/Stx2H6/E) which consists of two VHHs that each bind and neutralize Shiga toxin 2 (Stx2) (32). (**B**) Western blots of the VHH heterodimer following incubation (150 µg/ml) with a rabbit intestinal extract (1:20) in which the digestion was quenched after the indicated times of 37°C incubation. 10 µl replicate samples were resolved by SDS-PAGE, transferred to filters and then probed with one of two different rabbit serum pools, anti-denatured VHH sera or anti-native VHH sera, as indicated. Detection used HRP-coupled anti-rabbit Ig reagent. The band that we hypothesize to represent one or both of the functional ‘core’ VHHs in the heterodimer is indicated by an arrow. (**C**) Stx2-binding ELISA on samples from 4B that were quenched at the indicated times of incubation with chyme. Stx2-bound proteins were detected with rabbit anti-denatured VHH sera.

While anti-VHH sera western blots proved very useful as a tool to monitor the <u>structural</u> consequences of intestinal protease exposure to VHH proteins, it is the <u>functional</u> consequences to these VHH proteins that most impacts their potential use in enteric disease therapeutic applications. To monitor target-binding VHH function, we performed binding ELISAs with the rabbit anti-denatured VHH sera to detect Stx2-bound VHH peptides using the same chyme incubation samples tested in **Fig 4B**. ELISA results employing rabbit anti-denatured VHH sera readily detected VHHs that had bound to Stx2 (**Fig 4C**), even in samples incubated overnight in chyme that had been reduced to a single visible band of ∼14 kDa on western blot. Note that the higher peak signal for untreated control in this and other similar ELISAs likely reflects recognition of additional flanking epitopes that are lost rapidly during chyme incubations as well as some reduction in agent affinities resulting from dimers being cleaved into monomer components. Based on the shift in the EC_50_ values of the treated samples compared to untreated protein, it appears that at least 10% of the original Stx2 binding activity remained detectable by rabbit anti-denatured VHH sera following the overnight incubation with intestinal proteases under the *in vitro* conditions employed.

### Monomeric VHH agents vary widely in functional stability to intestinal proteases

Studies described in the prior sections found that exposure to enteric proteases rapidly removes epitopic tags of VHH multimers and cleaves spacers resulting in monomeric core VHHs which, on longer incubations, generally lose their function and integrity. We next sought to test whether different VHH monomers have similar or widely variable susceptibility to functional inactivation by more harsh exposures to intestinal proteases. For this, we took advantage of our prior discovery of four unrelated VHHs that all recognize the same small target of Stx2 B-subunit, and all potently neutralize Stx2-induced toxicity in HeLa cells. These VHHs were each subjected to varying times of exposure to a more concentrated rabbit intestinal chyme extract (1:5 dilution). At each time point, VHHs were quenched with a protease inhibitor cocktail and then later assayed for Stx2-neutralization potency in a cell-based assay (top row, **Fig 5A**). MTT assays assessed the amount of HeLa cell metabolic activity that remains following exposure to Stx2 toxin. Increased cell viability indicated the level of toxin neutralization that remained in each antibody sample dilution. The ability of antibody samples to protect cells from toxin despite greater amounts of sample dilution reflects increased antibody potency. The results demonstrated a wide range of functional chyme susceptibility in the four VHHs. The JEN-D10 VHH appeared the most susceptible to inactivation as no detectable Stx2-neutralizing activity was found after only 20 minutes of incubation. In contrast, JSY-F12 VHH appeared to be the most protease resistant VHH as the Stx-2 neutralizing potency was strong after 3 hours incubation and remained easily detected even after overnight incubation. The remaining two VHHs, JFG-H6 and JGH-G1 remained active after 1 hour or 3 hours incubation respectively, but both had lost function after overnight incubation. The same samples were also assayed for their functional integrity by western blot using the rabbit anti-denatured VHH sera for detection (**Fig 5B**). The results of the cell-based assay and western blots were highly consistent as the ∼14 kDa core VHH was detected only in the samples that displayed Stx2-neutralizing activity. These results also seem to confirm the interpretation that the 14 kDa bands, which display high resistance to intestinal protease digestion as described above, represent the highly structured core VHH.

**Fig 5.**
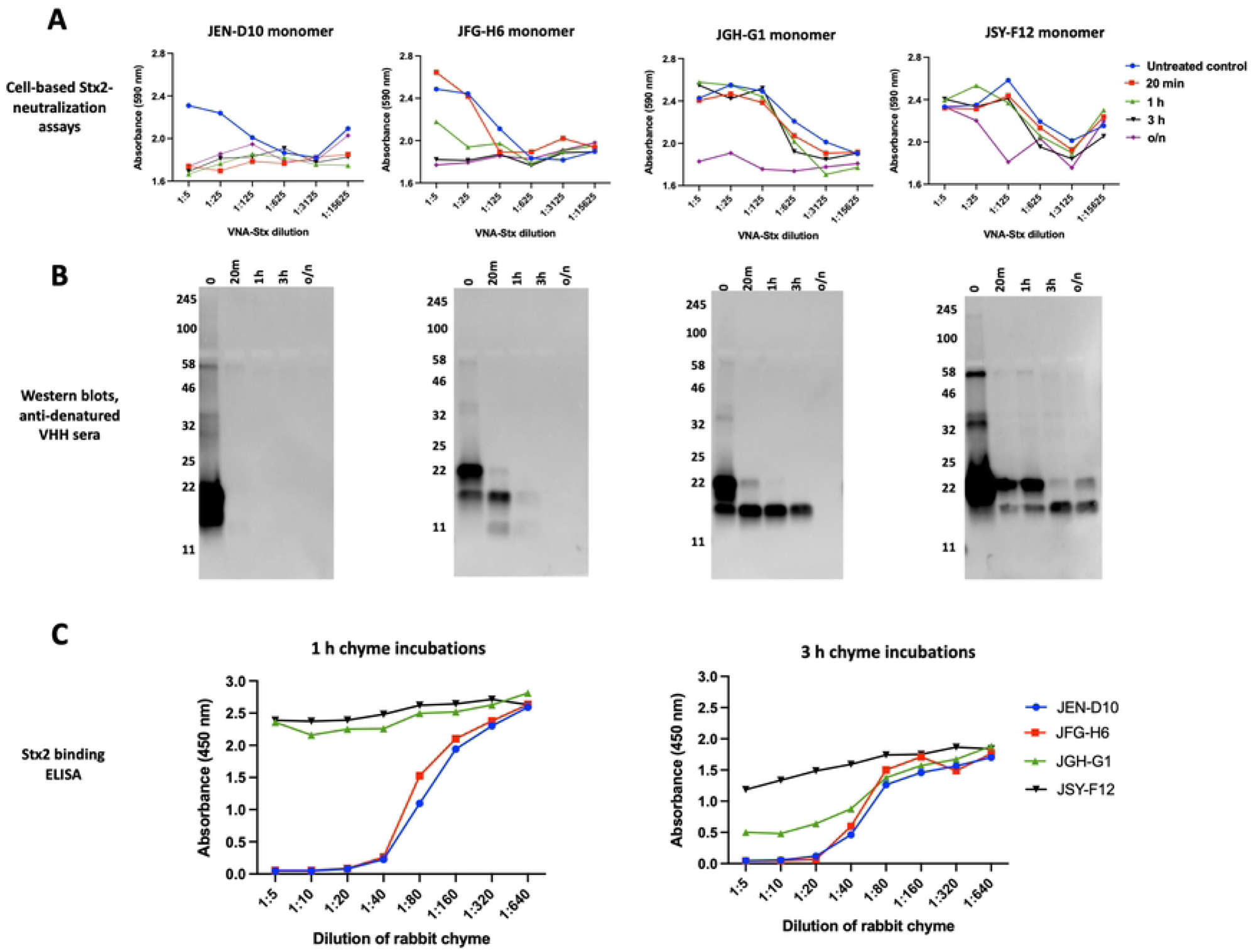
Monomeric VHHs vary widely in functional stability to intestinal proteases. (**A**) Functional stability of different Stx2-neutralizing VHH monomers following intestinal protease incubation. Stx2-neutralizing VHH monomers vary substantially in their susceptibility to loss of neutralizing activity from intestinal protease exposures. Four unrelated, Stx2-neutralizing VHHs expressed as Trx-fusion proteins were incubated (75 µg/ml) with rabbit chyme (1:3 chyme dilution) on ice. At indicated times, aliquots were diluted 1:5 into 1x HALT and rapidly frozen. Later the samples were assayed (MTT) for Stx2-neutralizing activity in HeLa cell-based viability assays using a 5-fold-dilution series of the treated VNA-Stx2 samples starting at 1:5 (v:v). **(B)** Structural integrity of different Stx2-neutralizing VHH monomers following intestinal protease incubation. The same rabbit chyme treated samples created for 5A were diluted 1:5 in 1x SDS sample buffer and 10 µl aliquots were assayed for VHH integrity by western blotting with anti-denatured VHH sera. **(C)** Rapid, high-throughput assay to assess the stability of VHH agents to retain target-binding function following incubations with an intestinal protease extract dilution series. Four Stx2-neutralizing VHHs were coated on wells (10 µg/ml) and incubated for 3 hours at 37°C with a 2-fold dilution series of porcine chyme, starting at 1:5 in PBS (v:v). After washing, an Stx2 binding ELISA was performed by incubating all wells with 0.5 µg/ml Stx2B/myc and detecting bound target with anti-myc mAb 9E10.

Functional assays such as the enzyme inhibition or toxin neutralizing assays shown in **Fig 2C** and **Fig 5A** are often challenging to perform in the presence of enteric proteases. We thus developed a rapid and more general enteric PK assay that assayed VHH-agent sensitivity to the loss of their target-binding activity during exposures to intestinal chyme extracts. To avoid variations in the immunodetection sensitivity to different VHH agents, we employed a sandwich ELISA in which coated VHHs were exposed to an intestinal chyme dilution series for either 1 hour or 3 hours before being washed and then incubated with the VHH target antigens. Successful VHH-capture of the target was then quantified immunologically using specific anti-target reagents. We find that retention of target binding by the coated VHHs to higher concentrations of chyme provides a simple measure of their functional stability to chyme exposures. For example, **Fig 5C** shows that the relative stability of the four monomer Stx2-neutralizing VHHs to chyme exposures when assayed by this simple method closely matched the results obtained using the more challenging Stx2-neutralization assays (**Fig 5A**).

### Comparing the functional stability of IgG1, IgA2 and VHH-based agents to intestinal proteases

It is not possible to exactly compare the functional stability to enteric proteases of conventional vs VHH-based antibodies due to the fundamental differences in structure of the two antibody types and the wide variation in chyme sensitivity of different VHHs. In a first effort to generally compare the protease stability of conventional and VHH antibodies, we tested the functional protease stability of an Stx2-neutralizing VHH heterodimer with that of an Stx2-neutralizing IgG1 (5C12) and a recombinant IgA2 that was engineered with the 5C12 V_H_ and V_L_ domains and tested both with or without the polymeric immunoglobulin receptor (pIgR). Parallel incubations with intestinal chyme at 1:5 and 1:3 chyme dilution were performed on the four different Stx2-neutralizing antibody types. All the samples were then assayed in a 1:5 dilution series for their Stx2-neutralizing potencies using the HeLa cell-based viability assay. The results showed substantial chyme protease resistance by IgG1 and IgA2 (+/- pIgR) in 1:5 chyme dilution even after overnight incubations (**Fig 6A top row**). Under the same conditions, the Stx2-neutralizing VHH heterodimer (JFG-H6/JGH-G1) was substantially less resistant to enteric proteases than the conventional antibodies, remaining active after 3 hours incubation and losing most activity during overnight incubations. Under the harsher 1:3 chyme incubation conditions, the Stx2-neutralizing activity of the IgG and IgA was somewhat lower after overnight incubation while the VHH heterodimer lost most of its potency after only 3 hours incubation (**Fig 6A bottom row**). The functional PK of the heterodimer thus approximately matched that of the JGH-G1 monomer, from **Fig 5A**. The results suggest that if we had selected more protease resistant component VHHs, such as JSY-F12 (**Fig 5A**), the VHH-based agents would have a functional chyme stability that is more similar to that of conventional antibodies.

**Fig 6.**
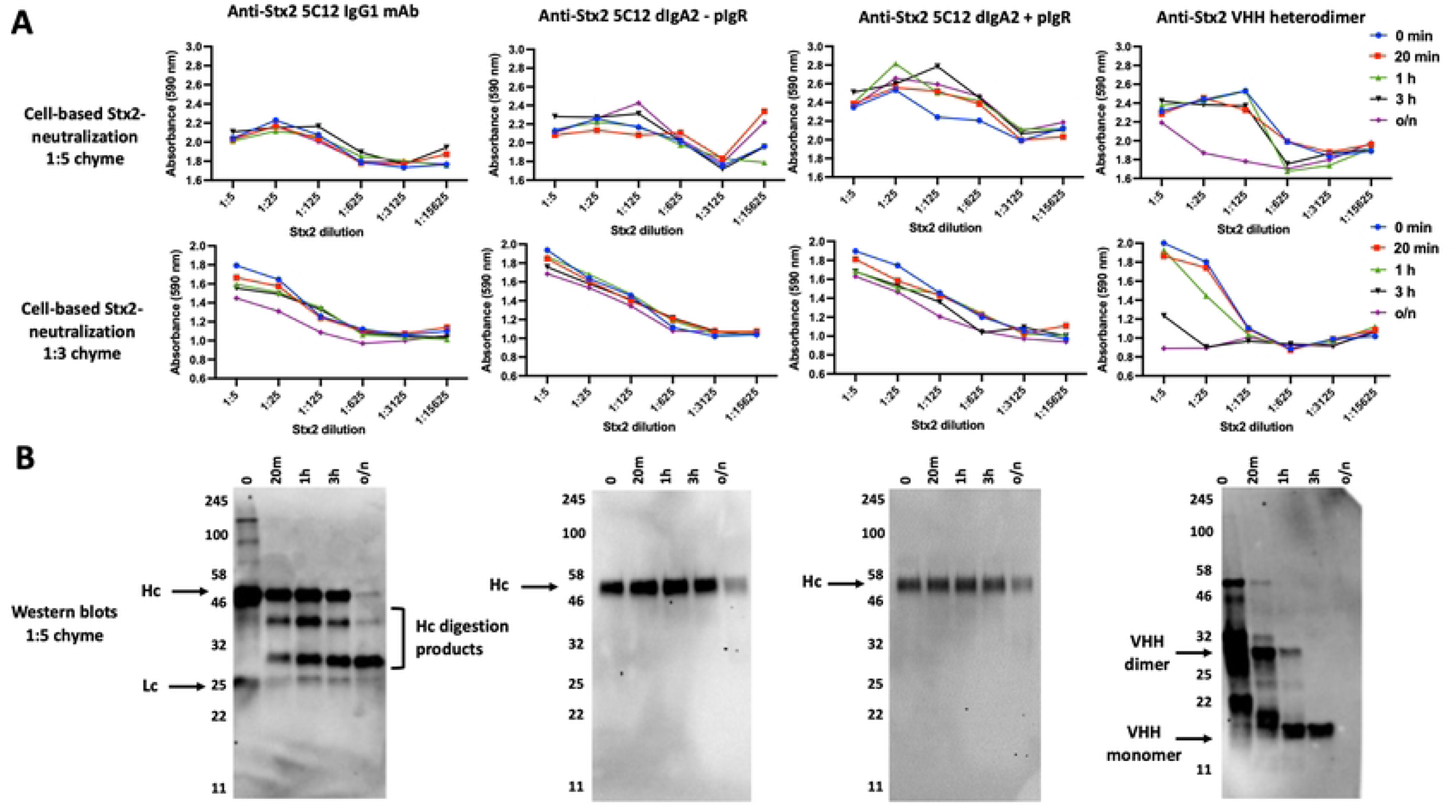
**Comparing the enteric PK of conventional antibodies IgG1 and IgA2 to that of VHHs**. (**A**) Stability of the Stx2-neutralizing potency of conventional and VHH-based antibodies following incubations with intestinal chyme. Stx2-neutralizing IgG1 mAb, 5C12 (300 µg/ml), or recombinant IgA2 engineered with 5C12 variable region 600 µg/ml (+/- pIgR, 400 µg/ml), or an Stx2-neutralizing VHH heterodimer (JFG-H6/JGH-G1, 100 µg/ml)) were incubated with rabbit chyme (1:5 or 1:3 chyme dilution as indicated) on ice. Samples were quenched at various times by 1:5 dilution into 1x HALT and then rapidly frozen. Later the samples were assayed for Stx2-neutralizing activity in HeLa cell-based viability assays with 5-fold-dilution series of the treated samples starting at 1:5 (v:v). **(B)** VHH integrity varies widely between different VHHs that were similarly exposed to intestinal proteases in a time course. The same chyme treated samples employed in 6A above were diluted 1:5 in 1x SDS sample buffer and 10 µl aliquots were assayed for antibody integrity by western blotting. The VHH agents and products were detected with rabbit anti-denatured VHH sera. To detect the IgG1 and degradation products, a goat anti-human IgG (H+L) was used. For IgA2 and heavy chain degradation products, a goat anti-human IgA (α chain) serum was used. Bands corresponding to heavy chain (Hc), light chain (Lc) and VHH dimer and monomer are indicated with arrows.

The same antibody samples assayed for Stx2-neutralization (**Fig 6A**) were also assessed for structural integrity by western blots (**Fig 6B**). The results revealed that the incubation time-dependent loss of function in **Fig 6A** showed similar kinetics to the loss of integrity of these antibody types. The IgG Hc appeared to be much more sensitive to degradation compared to the IgA2 Hc, which was not reflected in a loss of neutralizing potency, suggesting that loss of IgG1 Hc integrity did not significantly impact antibody potency to neutralize Stx2. This is not surprising as long as the Fab region remained intact as suggested by the western blot. As expected, the VHH heterodimer was rapidly degraded to VHH monomers by enteric proteases while the monomer integrity appeared to correlate well with the functional potency of the different incubation time points.

### Comparing VHH heterodimer spacer lengths and sequences for susceptibility to enteric proteases

Studies from our lab and others have demonstrated that VHH multimers generally exhibit higher affinities and greater potencies than VHH monomers (1). Unfortunately, as demonstrated above, VHH multimers are rapidly degraded by intestinal proteases within conventional spacers separating the VHH components. We attempted a range of studies to determine whether variations in the spacer size or sequence would prolong the functional stability of VHH heterodimers. Functional studies proved of limited value because the resulting monomers retain substantial potency and could not be sensitively distinguished from undigested multimers. Thus, western blots were more effective as a tool to monitor cleavage of the spacer regions separating VHHs.

**Fig 7** shows a study comparing the intestinal protease sensitivity of three anti-*Clostridium difficile* toxin B (TcdB) heterodimers containing TcdB-inhibiting VHH components 5D and E3 (**Fig 7A**). Two variables were introduced into the three heterodimer constructions, specifically the different spacers and/or flanking sequence modifications as detailed in **Table S1**. In this study, two of the heterodimers completely lacked a spacer between the two VHHs. To engineer this, the carboxyl end of the 5D core VHH (…VSS) was joined directly to the amino end of E3 core VHH (QVQLVE…). One of the two heterodimers lacking a spacer was flanked by two copies of a 10 amino acid peptide PEPEPEPEPE (PE5). The PE5 peptides were included to test whether flanking proline-rich peptides might protect the amino and carboxyl ends of the functional VHH heterodimer from degradation by amino peptidases and carboxypeptidases likely present in the chyme (17). In addition, proline/glutamic acid multimers were shown to form rigid peptides (18, 19) which could prove more resistant to intestinal protease cleavage. A third construction contained a more conventional flexible spacer consisting of 15 glycine residues (15G), in which the glycines were joined to the same core VHH sequences of 5D and E3, and also contained the proline-rich flanking sequences, The functional potency of the three heterodimers was tested following various times of incubation with rabbit chyme (1:5) in a Vero cell-based TcdB-neutralization assay, where cell rounding and detachment were measured to evaluate the extent of toxic effects from TcdB. As can be seen in **Fig 7B**, there were no apparent differences in the rate at which the three constructs lost their TcdB-neutralizing potency with increasing chyme incubation times. The same incubations were also analyzed by western blot employing the rabbit anti-denatured VHH sera and the results (**Fig 7C**) revealed no apparent differences in the rates at which the VHH heterodimers were degraded to monomers.

**Fig 7.**
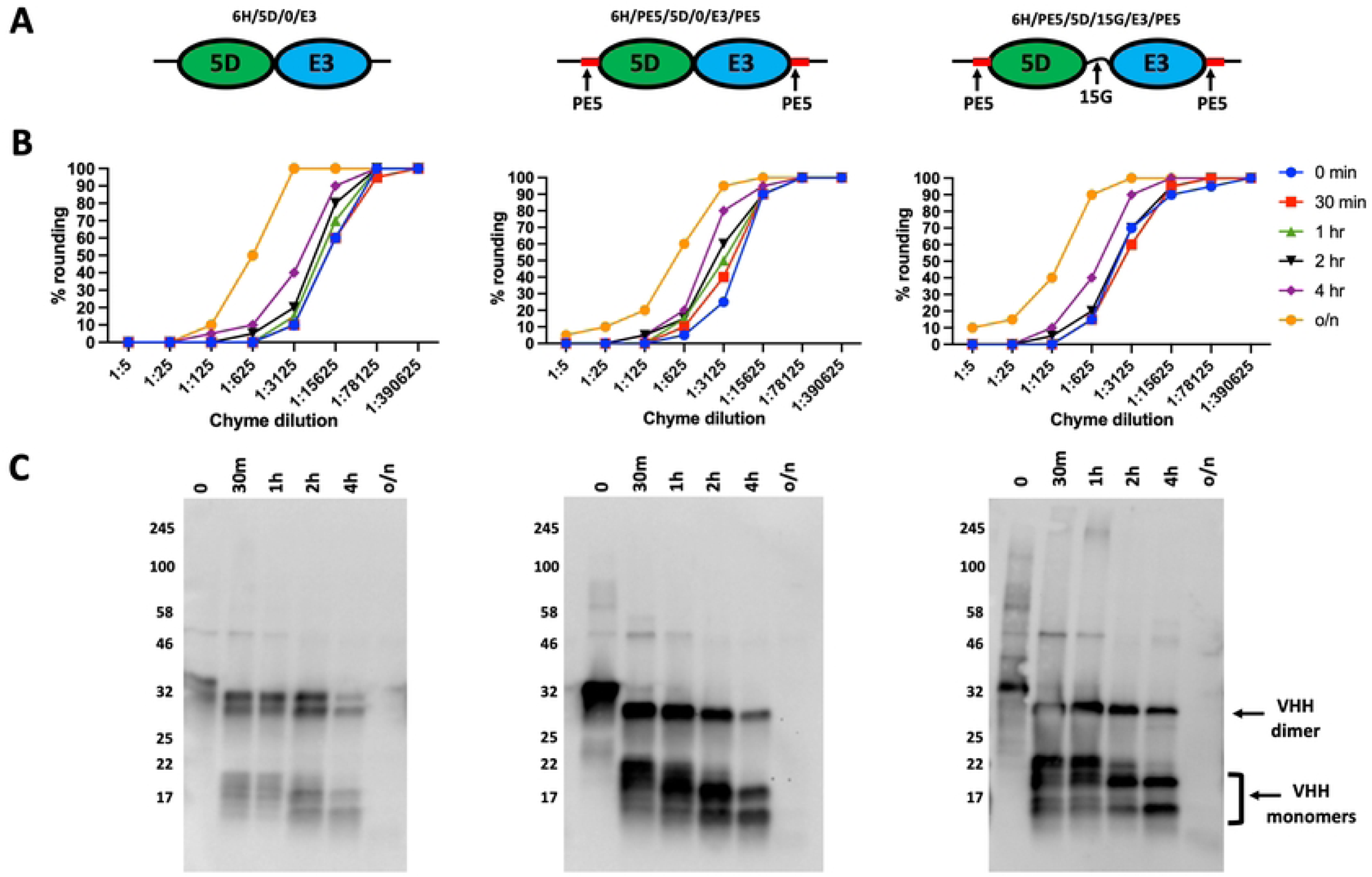
Enteric proteases similarly cleave VHH dimers with variable flanking sequence and spacer length. (**A**) Diagrams of VHHs used to test the enteric protease resistance of different VHH spacers and flanking sequences as shown in **Table S1**. Three VHH heterodimers, each consisting of two TcdB-neutralizing core VHHs, 5D and E3 (33), were engineered, expressed and purified as shown in diagram form. Each heterodimer was identical except that 6H/5D/0/E3 and 6H/PE5/5D/0/E3/PE5 had no spacer separating the VHH cores, and 6H/PE5/5D/0/E3/PE5 and 6H/PE5/5D/15/E3/PE5 each had the 10 amino acid sequence, PEPEPEPEPE (PE5) precisely flanking the pair of VHH cores. All had an amino end hexahistidine for purification and a carboxyl end myc tag. (**B**) The three VHH heterodimers shown in A (50 µg/ml) were each subjected to incubations with rabbit chyme (1:5) for the indicated times and quenched on ice with 1x HALT protease inhibitor cocktail. (**C**) The resulting samples were diluted 1:1 with 2x SDS sample buffer and were then assayed for their integrity by western blot (10 µl loaded per lane) and probed with rabbit anti-denatured VHH sera. Assay results are positioned below the diagrams of the VHH heterodimers being tested by each method.

In another study, we compared the intestinal protease resistance of three different VHH spacers expected to differ in their flexibility (**Fig S8a**). We employed the same VHH heterodimer in which the only difference in the three proteins was in the sequence of spacers separating the two VHHs as detailed in **Table S2**. One spacer consisted of a conventional 5 amino acid polyglycine flexible spacer (5G) while the other spacers were either the proline-rich amino acid rigid spacer, PEPEPE (PE3) or a proline/glycine hybrid spacer, PGPGPG (PG3). The three heterodimers were incubated at various times in rabbit chyme (1:3) and analyzed by western blot with rabbit anti-denatured VHH sera (**Fig S8b**). Again, we were unable to detect any meaningful difference in the rates at which the heterodimers were reduced to monomers.

### In vivo assays of VHH-agent enteric PK

In an effort to test the *in vivo* relevance of *in vitro* enteric PK studies using chyme incubations, we performed incubations of VHH-agents injection into ligated segments of extruded intestinal ‘gut loops’ as shown in **Fig 8A**. Three successive, linked duodenal gut loops were selected. Two loops were injected with an anti-TcdB heterodimer (cartoon shown in **Fig. 8B**) while a control loop located between them was injected with PBS to control for potential leakage (n=1). One sample was removed and quenched immediately after injection and massaging to mix the sample within each loop. Additional samples were removed at 15 min, 45 min and 2 hrs following loop injections. Functional *in vivo* persistence of the VNA-TcdB function during intestinal gut loop chyme incubation was assessed using a cell-based TcdB-neutralization cell rounding assay of the samples removed at different time points. The control gut loop sample displayed no significant TcdB neutralization (**Fig 8C, left panel**) showing that the chyme itself had negligible effect on the Vero cell rounding assay and that no detectable VNA-TcdB had passed through the surgical ties separating it from the adjacent gut loops injected with VNA-TcdB. As shown in **Fig 8C (middle and right panel)**, the gut loop samples obtained from the two loops injected with VNA-TcdB displayed clear TcdB neutralization at the earliest timepoint tested (1 min), preventing TcdB-elicited cell rounding even when diluted more than 1000-fold. The 1 min timepoint would have experienced some exposure to proteases prior to removal and quenching but provided an approximation of the initial potency of the VNA-TcdB diluted into the gut loop chyme. During the 2 hr *in vivo* gut loop incubation, the VHH heterodimer potency dropped about 5 and 20-fold in the replicate loops based on their estimated IC_50_s as compared to the 1 min timepoint. As a final measure for comparison, ELISAs were performed on the samples removed from VNA-TcdB injected loop (n=1) (**Fig 8D**) using rabbit anti-denatured VHH sera. As observed with the TcdB-neutralization assay, ELISA EC_50_ shifts indicated that the TcdB-binding potency of the samples dropped about 20-fold during the 2 hr *in vivo* gut loop residence. The results are within the range expected based on similar studies performed *in vitro* with chyme and support the value of using much simpler and more robust *in vitro* assays to obtain accurate estimates of the *in vivo* enteric functional PK of VHH-based agent samples. In addition, they indicate that target-binding ELISAs detected using rabbit anti-denatured VHH sera can provide a simpler and useful measure of the enteric functional stability of VHH-based agents.

**Fig 8.**
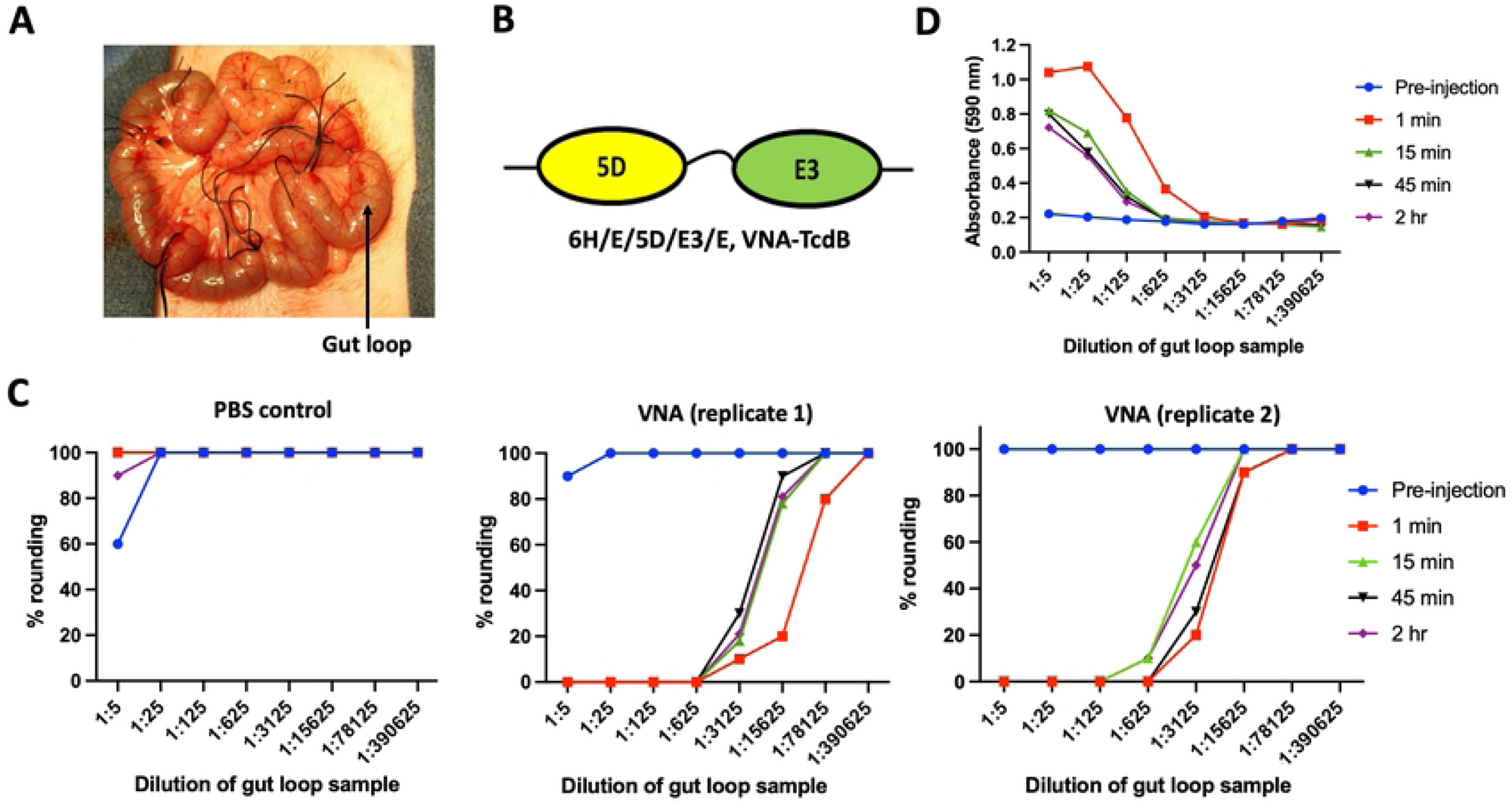
Comparing *in vitro* vs *in vivo* assays of VHH-agent enteric PK. (**A**) Photograph of gut loops extruded from anesthetized rabbit. The jejunum of an anesthetized rabbit was extruded through an incision and ligations were performed to tie off the intestine at several sites about 2 inches apart to create gut loops as shown. (**B**) Diagram of a VHH heterodimer (6H/E/5D/E3/E) which consists of two VHHs that each bind and neutralize *C. difficile* toxin B (TcdB) (33). The recombinant protein lacked the Trx fusion partner but retains an amino terminal hexahistidine (6H) and flanking E-tags. (C) Functional stability of VNA-TcdB to *in vivo* incubations within injected live gut loops. One gut loop was injected with 1 ml PBS and the two flanking gut loops were each injected with 1 ml PBS containing 500 µg of VNA-TcdB. After injection and mixing, 300 µl aliquots were removed at the indicated time points and immediately frozen in dry ice. Later the samples were thawed, diluted in protease inhibitors, filter-sterilized and assayed for TcdB inhibition potency. TcdB-neutralization assay results are shown for samples removed from the PBS injection control loop (C, left panel) and each of the two VNA-TcdB-injected loops (C, middle and right panels). (**D**) Functional stability of VNA-TcdB based on TcdB binding dilution ELISA on samples from a VNA-TcdB-injected gut loop. ELISA plates were coated with TcdB and VHH-agent products binding TcdB were detected with rabbit anti-denatured VHH sera.

## Discussion

Single-domain antibodies, such as VHHs, are particularly promising as components of enteric therapeutics because they can be produced at low cost in microbes and delivered orally or expressed in patients by probiotics or, possibly by engineering their own microbiome. Such products have already been reported to show some therapeutic benefits in numerous animal models of enteric diseases (2–11). Because the GI tract is a hostile environment for protein therapeutics due to its role in proteolytic food digestion and its rapid luminal flow leading to elimination, optimal exploitation of VHH-based agents as enteric therapeutics will require a good understanding of the factors that influence the functional stability of these agents while resident within this environment. We report here on some initial enteric PK studies with VHH-based agents.

Unsurprisingly, this hostile proteolytic environment of the GI tract produces substantial challenges to the performance of enteric PK studies. Initial PK studies, such as we report in **Fig 2** above, showed that conventional immunologic detection tools that employ epitopic tags fused to the VHH-based agents are of limited value as intestinal protease incubations quickly degraded the tags well before the VHH biological functions were lost. Clearly the functional stability of the VHH agents is a much more relevant property to monitor when performing enteric PK studies, yet these assays can also be difficult to perform quantitatively when the samples are contaminated with the highly variable, complex mix of components, including microbes and proteases, found within intestinal chyme.

For ideal PK studies, the intestinal protease components in chyme, and their concentrations, would be consistent and well known so as to permit the use of defined mixtures for *in vitro* enteric PK studies that mimic those in extracts of intestinal fluid. But we lack complete knowledge as to the identity and concentrations of each protease found in chyme and which proteases elicit the greatest influence on VHH-agent stability. It is known that pancreatic enzymes are stimulated by feeding (20), so large variations in enteric protease levels occur naturally, and also may vary substantially between different individuals, species and life stages. For all these reasons, we chose to employ crude extracts of chyme or feces from three different species for our initial studies as an approximation of the intestinal environment. Using enteric extracts for *in vitro* PK studies is clearly imperfect as it does not mimic the dynamic, flowing nature of the GI tract. Another complication of using crude enteric extracts is that incubations cannot be unambiguously terminated by addition of protease inhibitor cocktails due to the possible activity of unknown proteases that are not completely inhibited. If some proteases remain active, further degradation may occur during later functional assays leading to underestimates of the inherent functional stability of some agents. Despite these challenges, we were able to show that the PK results obtained from *in vitro* time course incubations of a VHH heterodimer with intestinal extracts compared favorably to similar time courses obtained *in vivo* by injecting the same VHH heterodimer into surgically extruded gut loops (**Fig 8**). In summary, we feel that the results reported in these studies are qualitatively significant, though quantitative outcomes must be considered as approximations.

Because the use of epitopic tags proved to be effective only when very brief or diluted chyme incubations were employed, we sought to identify or create an anti-VHH reagent better able to detect VHH agents that had been exposed to chyme for longer time periods. Preliminary studies with some commercial anti-VHH agents proved unproductive, so we created custom sera by immunizing two pairs of rabbits with a pool of VHH monomers. In one pair of rabbits, we employed native VHH immunogens in aqueous alum adjuvant to produce ‘anti-native VHH sera. In the second pair of rabbits, half of the VHH pool was denatured by boiling under reducing conditions prior to immunization and formulated in Freund’s oil-based adjuvant resulting in ‘anti-denatured VHH sera’. The same VHH monomer immunogens were included in all immunized rabbits, and shared small vector-derived peptides flanking the VHHs consisting mostly of epitopic and purification tags. Both rabbit sera pools strongly recognized intact VHH monomers on ELISAs and western blots (**Fig 4**). When testing the anti-native VHH sera, we found that virtually all of the VHH immunodetection signal rapidly lost at early times of chyme incubations, well before significant drops in VHH function were detected (**Fig 4B**), and similar to our observations of the epitopic tag sensitivity to chyme (**Fig 2B**). It was only when using rabbit anti-denatured VHH sera that we observed significant VHH recognition of samples incubated for longer times with high concentrations of chyme (**Fig 4B**). The presence of a 14 kDa product on western blots at later times of VHH agent chyme incubations appeared to correlate well with the retention of VHH function. We conclude that this band represents functional ‘core’ VHHs. Note that some VHH samples extensively degraded by chyme, such that VHH monomers were not detected by rabbit anti-denatured VHH sera, still retained detectable function in some cases (e.g., overnight treatments, **Fig 7B vs 7C**), indicating that there are limits to the immunodetection of bioactive VHHs even with this reagent. The full set of data reported here highlights the substantial challenges in performing quantitative enteric PK studies of VHH-based agents when employing immunodetection tools.

Of particular note, the ∼14 kDa core VHH chyme digestion product readily detected by anti-denatured VHH sera was virtually unrecognized by anti-native VHH sera. We believe these data provide additional evidence beyond that previously reported (21, 22), that the minimal functional core VHH itself is poorly immunogenic. Furthermore, this finding indicates the importance of omitting foreign peptides flanking the core VHH components within VHH-based therapeutics when anti-drug antibody responses need to be avoided.

Our results show that there are significant challenges to the use of VHH multimers or VHH fusion proteins as enteric therapeutics for applications requiring the components to remain connected while in the GI tract. The most sensitive sites of intestinal protease cleavage of VHH multimers were determined for several heterodimers (**Fig 3**) and each occurred a few residues to the amino end of a core VHH. It is difficult to know whether the small amino acids (A, G, Q) surrounding the cleavage sites reflect a preferred protease specificity or simply occurred at sterically available sites immediately flanking the structural core VHH. We thus sought to proactively engineer spacers and flanking sequences into VHH-based agents in efforts to slow the intestinal protease activity. Numerous different spacers and flanking sequences were tested yet all remained highly susceptible to cleavage by intestinal extract incubations at times before the VHH functions themselves were significantly damaged. In one approach we tested the use of rigid or flexible spacers and exoprotease-resistant flanking peptides (**Fig 7, S8**). We also tested core VHHs that were linked together without any intervening spacer, hypothesizing that removal of this small region of protease-susceptible unstructured peptide would resist cleavage by the enteric proteases, however even this minimal linkage remained protease susceptible (**Fig 7**). Additional spacer sequences were less rigorously tested in other studies not reported here, and again none displayed measurably improved protease resistance. In total, the data seems to suggest that the intestinal proteases rapidly degrade unstructured peptides such as epitopic tags and spacers and are much slower to degrade highly structured proteins such as the core VHHs. Since most proteins from food entering through the stomach are denatured by acidic conditions, it makes sense that the intestinal proteases would have their highest specificity for unstructured protein.

Our unsuccessful efforts to identify spacers that prevent VHH multimers from being cleaved to monomers, even when the VHH cores were directly joined, bodes poorly for the ability to maintain VHH multimers or fusions to partners in the GI tract. Thus, in enteric situations when VHH monomers are not sufficient, it may prove necessary to employ alternative approaches in which the VHH fusion proteins are continuously expressed and thus replenished within the GI tract such as by living probiotic bacteria. This strategy has been successfully demonstrated recently in two systems; one expressing VHH/CsgA-fusions within probiotic *E. coli* Nissle which produced VHHs as stacked multimers within curli fibers (11), and another displaying VHHs fused to a surface protein of *Lactobacillus paracasei* within the GI tract (23).

Despite the substantial resistance of core VHH monomers to proteolytic inactivation by intestinal extracts, the functional activity of all tested monomer VHHs was eventually destroyed with prolonged chyme exposure. We performed a series of studies comparing the functional stability of individual monomer VHHs to incubations with intestinal extracts. In one study, we compared the functional stability of a panel of four unrelated Stx2-neutralizing VHH monomers (**Fig 5A, 5B**). The results revealed dramatically different enteric PK properties of the four VHH monomers assessed under the same treatment conditions. Since the success of VHH-based products as enteric therapeutics is likely to depend on the use of VHHs that are highly resistant to destruction by enteric proteases, we developed and validated a rapid and general method to assess enteric PK properties of monomeric VHHs by sandwich ELISA (**Fig 5C**). Anecdotally, our lab has performed similar rapid assays on dozens of other VHHs having a wide variety of target specificities as a down-selection screen for VHHs that show high functional resistance to enteric proteases. To detect VHH target binding in these assays, we typically use the immune alpaca antisera generated previously for the original VHH discovery. These studies have continued to reveal wide variations between different VHHs in their functional stability to intestinal chyme.

Finally, we performed preliminary studies seeking to generally compare the enteric PK of VHH agents with conventional IgG and IgA antibodies (**Fig 6**). Because the target binding domains of mAbs and VHHs are inherently different, the comparisons were at best approximations. The results showed that the toxin-neutralizing functions of the IgG and IgA2 were more stable to incubations with intestinal extracts than the VHH-based agent we tested. Both components of the tested VHH dimer, though, were only moderately resistant to enteric proteases (**Fig 5**) and our results suggest that use of VHHs that are pre-selected for high protease resistance will permit development of VHH-based products with enteric PK properties more similar to that of conventional antibodies.

In summary, we report studies investigating the enteric PK of VHH-based agents in an effort to provide information to facilitate the exploitation of these unique agents as components in the development of immunotherapeutic agents with potential to treat enteric diseases. The findings show that while VHH monomers vary widely in their functional stability to the proteolytic environment of the GI tract, a subset of VHHs possess a stability that is similar to that of IgG and IgA mAbs. The findings also reveal the substantial challenge of maintaining the integrity of VHH multimers and other proteins and peptides to which they are fused, including epitopic tags used in their detection. In total, this work should provide useful guidance to scientists that intend to use VHH nanobodies for any purpose in which they may be exposed to the enteric environment.

## Materials and Methods

### Ethics statement

All *in vivo* studies were performed within the guidelines established by the Guide for the Care and Use of Laboratory Animals of the National Institutes of Health and were approved by the Tufts University Institutional Animal Care and Use Committee (IACUC) under Protocols No. G2017-18, No. G2017-55 and No. G2017-119.

### Enteric extract preparations and incubations

Variable segments of the rabbit or porcine GI tract representing sections of duodenum, jejunum, ileum, cecum, colon and rectum were removed by dissection and the contents were squeezed out into 50 ml tubes and diluted approximately 1:4 (v:v) in PBS. Human fecal solids from volunteer donors were similarly diluted approximately 1:4 (v:v) in PBS. Tubes were then vigorously vortexed and centrifuged (∼10,000 g). The resulting chyme or fecal extract supernatants were stored in aliquots at -80°C as 1x chyme or 1x fecal extract. Different antibody preparations and controls were incubated at 37 °C with extracts of human feces or intestinal chyme of porcine or rabbit origin in PBS, and reactions terminated with either α1-antitrypsin (α1AT) (Sigma) or 1x HALT^TM^ protease inhibitor cocktail (Thermo Fisher), using the concentrations and conditions indicated.

### Gut loop preparation and incubations

Specific pathogen-free New Zealand White rabbits (Covance) were anesthetized with ketamine (35mg/kg intramuscularly (24)) and xylazine (5mg/kg IM) and an intravenous (IV) catheter and endotracheal tube were placed. Anesthesia was maintained with isoflurane, 1-2% by inhalation. Respiratory rate, oxygen saturation (SpO_2_; %), and end tidal CO_2_ were monitored and normocapnia (35–45mmHg) was maintained. Rabbits were kept warm using a heating pad and lamp during the experiment, IV crystalloids were administered for maintenance fluid support, and rectal temperature and blood pressure (indirect oscillometric measurement from the forelimb) were continuously monitored. The ventral abdomen was clipped and prepped aseptically. Using sterile surgical technique, an 8 cm midline abdominal incision was made, and a segment of jejunum exteriorized. Adjacent jejunal segments (5–6 cm each) were isolated by circumferential ligation of the jejunum and the terminal arcade of the jejunal artery and vein, as described previously (25). Adjacent, isolated jejunal segments (approximately 6 cm each) were created using circumferential ligations (2-0 silk) around the intestine. Care was taken to avoid injury to the mesenteric vessels and not compromise the blood supply. Injection into an isolated jejunal segment initiated a study and sampling of the contents was then performed serially. The intestines were placed back into the abdominal cavity and the abdominal wall closed between each loop sampling to maintain body and abdominal temperature. The order of the treatment groups was randomized between rabbits.

### VHH and VHH multimer expression

VHHs and VHH heterodimers were expressed in *E. coli* as previously reported (26). In some cases, monomer VHHs were expressed in a vector deriving from pET32 which lacked the thioredoxin (Trx) fusion partner (Trx coding DNA was removed by Nde1 digestion and vector re-ligation) such that VHH products contained an amino terminal hexahistidine for purification and a carboxyl-end myc tag for detection. Expression of the heterohexameric protein, VNA1-ABE, was performed in mammalian CHO cells as previously described (13).

### Generation of IgG and IgA antibodies

The Stx2-neutralizing IgG mAb, 5C12, was produced as previously described (27). The IgA2 version of 5C12 was produced using previously described methods (24). Briefly, to generate expression constructs, variable-region sequences were ordered as gBlock DNA fragments (Integrated DNA Technologies) and NEBuilder HiFi DNA Assembly Master Mix (New England Biolabs) was used to clone these into the pcDNA3.4 (ThermoFisher) backbone plasmid as expression cassettes flanked on the 5’ end by the DNA encoding leader sequences from either murine variable kappa chain (ATGGGTGTGCCCACTCAGGTCCTGGGGTTGCTGCTGCTGTGGCTTACAGATGCCAGATGC) or a murine variable heavy chain (ATGGAATGGAGCTGGGTCTTTCTCTTCTTCCTGTCAGTAACTACAGGTGTCCACAGC); on the 3’ end, the constructs encoded the human kappa-chain constant region (IGKC*01; GenBank:J00241) or the human alpha-chain constant region (IGHA2*01; GenBank:J00221). Similarly, a gBlock encoding human J-chain (hIgJ) sequence (GenBank:XM_011531926) was cloned into the pcDNA3.4 backbone plasmid. Human polymeric immunoglobulin receptor (hpIgR) construct was assembled from a cDNA clone (Sino Biological, cat. HG10131-UT), which was used to extract the fragment encoding the hpIgR ectodomain (GenBank:NP_002635; amino acids 19–638), and was placed between DNA fragments encoding tissue plasminogen activator leader peptide (28) and a C-terminal octahistidine tag (8H) followed by an AviTag (29) into the pcDNA3.4 backbone plasmid, resulting in the hIgR.HisAvi plasmid.

For the dimeric IgA production, FreeStyle 293-F (Thermo Fisher) cells were transiently co-transfected using PEI MAX (Polysciences) with the 5C12 hIgA2/hIgK/hIgJ plasmids, using the following ratio of constructs 0.25/0.25/0.5. Following 5 days of culturing, conditioned medium was harvested by centrifugation, supplemented by the addition of NaN_3_ (0.02% final concentration), and NaCl (350 mM final concentration). Protein was then captured using Pierce Protein L Plus Agarose (Thermo Fisher), washed with HBS-E-hs buffer (10 mM HEPES, pH 7, 300 mM NaCl, 2 mM EDTA), and eluted in 0.1 mM glycine, pH 2.7 (fractions were immediately pH-neutralized using 1 M Na_2_HPO_3_). Protein-containing fractions were pooled and buffer-exchanged against HBS-E (10 mM HEPES at pH 7, 150 mM NaCl, 2 mM EDTA) by ultrafiltration using Amicon Ultra devices (Millipore).

Recombinant human pIgR (hpIgR) was produced by transient transfection with the hpIgR.HisAvi construct. Following 5 days of culturing, conditioned medium was harvested by centrifugation, and supplemented by the addition of NaN_3_ (0.02% final concentration) and NaCl (350 mM final concentration). Protein was then purified by immobilized metal-affinity chromatography using HisPur Ni-NTA resin (Thermo Fisher) followed by gel-filtration over a HiLoad 16/600 Superdex 200 pg column (Thermo Fisher) in the HBS-E buffer (10 mM HEPES, pH 7, 150 mM NaCl, 2 mM EDTA).

### Rabbit anti-VHH serum

Two New Zealand white rabbits (Covance) were immunized with a pool containing equal amounts of thirteen random, unrelated VHH proteins, expressed as above with only hexahistidine and myc tags, and administered under native, aqueous conditions to minimize VHH denaturation. Each rabbit was injected with 200 µg of the VHH pool followed by four subcutaneous immunizations with 100 µg of the same VHH pool, all using an aqueous alum adjuvant. Serum was obtained from rabbits several days after the final immunizations from a terminal bleed using standard procedures. The resulting serum pool was called ‘rabbit anti-native VHH sera’.

A second pair of New Zealand white rabbits were immunized with the same pool of VHHs as above except that, prior to immunization, 50% of the pool was subjected to reduction and denaturation by adjusting the solution to 1 mM DTT and boiling for 30 minutes. The remaining 50% of the pool contained native VHHs. For the second pair of rabbits, the first three immunizations employed complete Freund’s adjuvant (CFA) while the last two employed alum adjuvant. This serum pool was called ‘rabbit anti-denatured VHH sera’.

### ELISAs and western blots

ELISAs and western blots were performed as previously described (30) with specific details provided in figure legends. Detection of VHH agents employed: 1:1000 rabbit anti-native, or anti-denatured VHH sera followed by goat HRP/anti-rabbit IgG (Sigma); rabbit HRP/anti-E-tag (Bethyl); or rabbit HRP/anti-GFP (Santa Cruz). Detection of mAb employed: goat HRP/anti-mouse IgG (Santa Cruz); donkey HRP/anti-human IgG (Jackson Research Labs), or goat HRP/anti-human IgA (Sigma). Detection of IgG1 employed anti-human IgG (V_H_+V_L_) (Jackson) and of IgA2 anti-human IgA (Hc only) (Sigma). Commercial antibodies were used at dilutions recommended by manufacturer. Signals were detected using peroxidase reagent and colorimetric or chemiluminescent substrate, and measured as described elsewhere (30).

### Sandwich ELISAs

The four Stx2-neutralizing VHHs were coated overnight at 10 µg/ml concentration onto individual columns of a 96-well Nunc Maxisorp ELISA plates, then washed but not blocked. A 2-fold dilution series of porcine intestinal chyme was applied starting at 1:5 in PBS in the top well and down the plate. The plates were then incubated at 37°C for either 1 hour or 3 hours before extensive washing and blocking of the plate overnight in 1xPBS/4% milk/0.1% Tween. An Stx2 binding ELISA was performed on each plate by incubating the plate in 0.5 µg/ml Stx2, washing and detecting bound Stx2 with an anti-Stx2 (A subunit) mAb (5C12 (31)).

### Amino acid sequencing of chyme incubated VHH-agents

VHH proteins were incubated with enteric proteases for indicated times and resolved on a 4–20% Novex™ Tris/Glycine SDS/PAGE gel (Invitrogen). The gel was then electroblotted onto PVDF membrane and visualized with 0.02% Coomassie Brilliant blue in 40% methanol, 5% acetic acid for 20-30 seconds. Bands of interest were excised and subjected to N-terminal sequence analysis on an ABI 494 Protein Sequencer by the Tufts University Core Facility. The resulting mixture of sequences was analyzed blind to the samples by experienced staff in the facility.

### Cell-based toxin neutralization assays

Assay test samples from *in vitro* and *in vivo* incubations with enteric protease extracts were stored at -80°C until use. Prior to assays, samples were thawed on ice and diluted 30-fold in ice cold PBS containing 1x HALT^TM^. Diluted samples were then sterile filtered by syringe keeping all samples on ice prior to use in cell-based assays.

#### Stx neutralization assay

Stx2 neutralization was assayed in a cell-based assay as described previously (32). In summary, HeLa cells (ATCC) were incubated with Stx2 +/-VHH sample dilutions and cell viability was assessed 48 hours later by MTT assay.

#### TcdB neutralization assay

TcdB neutralization was assayed in a cell-based assay as described previously (33). Specifically, Vero cells (ATCC) were incubated with TcdB +/- VHH sample dilutions and cell rounding in each well was later estimated by microscopic observation. The toxin dose used was chosen as the final concentration that elicits 100% cell rounding in cells after about 2 hrs. The readings of all wells were obtained at the post-exposure time at which control cells (negative control VHH) had become 100% rounded, typically after 2-4 hours.

#### LC/A inhibition assay

This assay was performed as previously described (14) using the BoTest A/E reporter (BioSentinel) as the substrate.

## Acknowledgements

We gratefully acknowledge the excellent assistance of Jean Mukherjee, Alexa Foss for immunizing and caring for the rabbits. We thank David Lee-Paritz for assistance with the rabbit gut loop work and Sue Chapman for technical assistance. We also thank Michael Berne at the Tufts University Core Facility (TUCF) for the Edman degradation and amino acid sequence analysis.

## Abbreviations

sdAb: single-domain antibody
VHH: heavy-chain-only antibody V_H_ domain
VNA: VHH-based neutralizing agent
PK: pharmacokinetics
GFP: green fluorescent protein
YFP: yellow fluorescent protein
CFP: cyan fluorescent protein

## Supplementary Materials

**Fig S1. LC/A inhibition assay data in support of** Fig 2. Supporting data for Fig 2. Western blots on the left derive from 15m or 60m incubations of LC/A and its substrate, BoTest^™^ A/E (BioSentinel Pharmaceuticals) (34), in which aliquots were added from one of the two indicated VHH heterodimer incubations with intestinal proteases. One VHH heterodimer was Trx/ALcH7/JPUA5/E, which is a potent inhibitor of LC/A. The second heterodimer is Trx/JLDG10/JLEE9/E is an irrelevant control binding to BoNT/E holotoxin. The VHH heterodimer samples loaded on the gel derive from 0m, 10m or 60m incubations with either PBS, human fecal extract (hFecal) or porcine cecum extract (pCecum). Incubations were quenched and loaded for SDS-PAGE. The gel was then transferred to filters which were probed with an anti-GFP reagent. The relative amount of uncleaved BoTest^™^ A/E substrate was assessed on an imaging system by comparing the signal to that of undigested substrate (0m, PBS). Samples able to neutralize LC/A inhibition after chyme incubations protect the substrate from LC/A digestion. The full-size blot from the 15m digestion is shown on the right.

**Fig S2. Edman amino terminal amino acid sequences of the VHH heterodimer Trx/E/AH3/AA6/E in support of** Fig 3. Coomassie-stained SDS-PAGE **(A)** and Coomassie-stained PVDF membrane **(B)** of samples that derive from chyme incubations of VHH heterodimer Trx/E/AH3/AA6/E (33) (diagram shown in **C**). This VHH agent was incubated for 60 minutes (60m) or overnight (o/n) with 1:10 pig intestinal extract or 1:10 human fecal extract before 5 µg of VHH agent was loaded to each lane. Untreated VHH (0) was utilized as a control. Replicate lanes of selected samples were resolved by SDS-PAGE and transferred to PVDF, then the filter was stained (B). The two ∼16 kDa digestion products indicated in B with an arrow were submitted for amino terminal Edman sequence analysis (**Fig S3A, B**).

**Fig S3. Amino acid analysis traces for VHH heterodimer Trx/E/AH3/AA6/E sequential Edman degradation of sample from Fig S2**. Edman degradation traces of VHH heterodimer Trx/E/AH3/AA6/E digestion products from **Fig S2** resulting from incubation with **(A)** porcine intestinal chyme or **(B)** human fecal extract. The five traces corresponding to the first five residues are shown with the top candidates for amino acids 1 to 5 indicated with arrows. The top amino acid calls (identified blind) are shown to the right, with secondary calls in parentheses. The sequencing results of both bands indicated an amino terminus of AQGVQ which could be unambiguously identified at the amino end of the submitted peptide shown in Fig 3 and **Fig S2C**.

**Fig S4. Edman amino terminal amino acid sequence of the VHH heterodimer Trx/E/JDQF12/JDQD12/E in support of** Fig 3. Coomassie-stained SDS-PAGE **(A)** and Coomassie-stained PVDF membrane **(B)** of samples that derive from chyme incubation of VHH heterodimer Trx/E/JDQF12/JDQD12/E (35) (diagram shown in **C**). This VHH agent was incubated for 10 (10m) or 60 minutes (60m) with 1:10 pig intestinal extract before 4 µg of VHH agent was loaded to each lane. Untreated VHH (0) was included as a control. Replicate lanes of selected samples were resolved by SDS-PAGE and transferred to PVDF, then the filter was stained (B). The ∼30 kDa digestion product indicated in B was submitted for amino terminal Edman sequence analysis (**Fig S5**).

**Fig S5. Amino acid analysis traces for VHH heterodimer Trx/E/JDQF12/JDQD12/E sequential Edman degradation of sample from Fig S4**. Edman degradation traces of VHH heterodimer Trx/E/JDQF12/JDQD12/E digestion products from **Fig S4**. The five traces corresponding to the first five residues are shown with the top candidates for amino acids 1 to 5 indicated with arrows. The top amino acid calls (identified blind) are shown to the left, with secondary calls in parentheses. The sequencing results indicated an amino terminus of AQVQL which could be unambiguously identified at the amino end of the submitted peptide shown in **S4C**.

**Fig S6. Edman amino terminal amino acid sequence of the VHH heterodimer 6H/E/StxA5/Stx2G1/E in support of** Fig 3**. (A)** Coomassie-stained PVDF membrane of samples that derive from chyme incubation of VHH heterodimer 6H/E/StxA5/Stx2G1/E (32) (diagram shown in **B**). This VHH agent was incubated with pig intestinal extract (1:30) and 2.5 µg VHH heteromultimer was then loaded per lane. After 60m, this ∼33 kDa VHH heterodimer resulted in a single digestion product of ∼14 kDa that likely consisted of both VHH components, each having similar amino terminal sequences, cleaved within the spacer. The band was excised as indicated in A and submitted for amino terminal Edman sequence analysis (**Fig S7**).

**Fig S7. Amino acid analysis traces for VHH heterodimer 6H/E/StxA5/Stx2G1/E sequential Edman degradation of sample from S6**. Edman degradation traces of VHH heterodimer 6H/E/StxA5/Stx2G1/E digestion products from **Fig S6**. The seven traces corresponding to the first seven residues are shown with the top candidates for amino acids 1 to 7 indicated with arrows. The top amino acid calls (identified blind) are shown at the top, with secondary calls in parentheses. We conclude that the correct amino end call for residue 1 must be a glycine as no similar sequence existed in this protein in which alanine could be the correct call. The sequencing results of the degradation product(s) indicated an amino terminus of GVQAQLQ, a sequence found immediately upstream of both core VHHs as shown in **Fig S6B**. We suggest that the submitted product was most likely a pool of two degradation products resulting from cleavages at these two sites in both VHHs. We cannot rule out that the band resulted from cleavage at just one of the two identified sites while the second VHH was further degraded to smaller fragments.

**Fig S8. Enteric proteases similarly cleave a VHH dimer varying in spacer sequences**. **(A)** Cartoons of the three VHH heterodimer proteins tested. Each heterodimer was identical except for the sequence of spacer region as indicated in **Table S2**. Each consisted of two Stx2-neutralizing core VHHs, JGH-G1 (G1) and JFG-H6 (H6) (32), an amino end hexahistidine for purification and a carboxyl end myc tag. 6H/G1/5G/H6 had a spacer consisting of 5 glycine residues (5G), 6H/G1/PG3/H6 had a spacer consisting of the six amino acid sequence PGPGPG (PG3), and 6H/G1/PE3/H6 had a spacer consisting of the six amino acid sequence PEPEPE (PE3). **(B)** The three VHH heterodimers (50 µg/ml) were each subjected to incubations with rabbit chyme (1:10) for the indicated times and quenched on ice with 1x HALT protease inhibitor cocktail. The resulting samples were diluted 1:1 with 2x SDS sample buffer and were then assayed for their integrity by western blot (10 µl loaded per lane) probed with rabbit anti-denatured VHH sera.

**Table S1. Flanking and spacer sequences of TcdB-neutralizing VHH heterodimer from Fig. 7.** Colored sequences correspond to VHH components diagramed in **Fig. 7A**.

**Table S2. Flanking and spacer sequences of two Stx2-neutralizing VHH heterodimer from Fig. S8.** Colored sequences correspond to VHH components diagramed in **Fig. S8A**.

## Notes

### Competing Interest Statement

The authors have declared no competing interest.

